# A Trisomy 21 Lung Cell Atlas

**DOI:** 10.1101/2023.03.30.534839

**Authors:** Soumyaroop Bhattacharya, Caroline Cherry, Gail Deutsch, Birth Defects Research Laboratory (BDRL), Ian A. Glass, Thomas J. Mariani, Denise Al Alam, Soula Danopoulos

## Abstract

Trisomy 21 (T21), resulting in Down Syndrome (DS), is the most prevalent chromosomal abnormality worldwide. While pulmonary disease is a major cause of morbidity and mortality in DS, the ontogeny of pulmonary complications remains poorly understood. We recently demonstrated that T21 lung anomalies, including airway branching and vascular lymphatic abnormalities, are initiated in utero. Here, we aimed to describe molecular changes at the single cell level in prenatal T21 lungs. Our results demonstrate differences in the proportion of cell populations and detail changes in gene expression at the time of initiation of histopathological abnormalities. Notably, we identify shifts in the distribution of alveolar epithelial progenitors, widespread induction of key extracellular matrix molecules in mesenchymal cells and hyper-activation of IFN signaling in endothelial cells. This single cell atlas of T21 lungs greatly expands our understanding of antecedents to pulmonary complications and should facilitate efforts to mitigate respiratory disease in DS.

---

Trisomy 21 (T21), resulting in Down Syndrome (DS), is the most prevalent chromosomal anomaly worldwide. The partial or complete triplication of human chromosome 21 (HSA21) results in numerous phenotypic abnormalities and complications, encompassing several organ systems. Although often associated with congenital heart disease (CHD), pulmonary complications and respiratory conditions remain one of the leading causes of hospitalization and mortality in children with DS 3 years of age or younger^1^. Several groups, including our own, have demonstrated that children can present with these pulmonary complications regardless of CHD^2–4^. Pulmonary complications in individuals with DS may span the entire lung, ranging from bronchomalacia in the large airways to alveolar simplification^2, 5^. Considering the prevalence and severity of these complications, the ontogeny of these defects warrants a better understanding.

Lung development is achieved through cross-talk of multiple, different cell lineages^6^. The anatomical and cellular structure of the fetal lung differs significantly from what is observed in the postnatal lung capable of gas exchange^7, 8^. We recently demonstrated that the developing T21 lung presents with histopathological abnormalities starting as early as 16 weeks of gestation^9^. Anomalies are observed in approximately 70% of those in the late pseudoglandular/early canalicular stage, with varying degrees of phenotypic severity. One of the pathways that is consistently hyper-activated in multiple T21 tissues is the type I Interferon (IFN) pathway^10^. In prenatal T21 lungs, we found evidence of IFN dysregulation at the time of initiation of histopathologic abnormalities associated with altered activation of both the complement and coagulation cascades and extracellular matrix (ECM) pathway^9^. Targeted studies indicating changes in cell proliferation and differentiation suggest cellular composition is altered in the developing T21 human lung^9^.

In this study we aimed to define changes in gene regulatory networks at the cellular level by performing single cell RNA sequencing on T21 lungs manifesting with developmental anomalies. Here, we report changes in the distribution of cellular phenotypes, and key gene expression alterations in discrete cell sub-populations in fetal T21 lungs. Additionally, we note that the pathways and cellular processes being altered in the different cellular lineages are sometimes overlapping, but often distinct, highlighting the complexity of impact from triplication of HSA21. This cellular atlas comprehensively defines regulatory processes associated with the initiation of pulmonary anomalies observed in T21 and may facilitate the identification of therapeutic targets for respiratory disease in DS individuals.

## Results

### Prenatal T21 lungs display all major cell lineages

To characterize cellular heterogeneity in the developing human fetal trisomy 21 (T21) lung as compared to age matched controls, we performed single cell RNA sequencing (scRNAseq) of manually and enzymatically dissociated cells from lungs 17-19 weeks of gestation. We previously established that histopathological anomalies are observed in T21 lungs as early as 16 weeks of gestation, with approximately 70% of lungs at this gestational age or older presenting with anomalies^9^. Seeking to better understand what could be contributing to these developmental defects, we sequenced 5 T21 lungs, each displaying some degree of histopathologic alteration (Supplemental Table S1), and 4 non-T21 control lungs. We analyzed a total of 36,259 cells (17,623 T21 cells and 18,636 non-T21 cells), with an average detection of 2,372 genes per cell. All the major cell lineages were identified, with the majority of cells being of mesenchymal origin (66.67%), followed by epithelial (16%), endothelial (9.77%), and lastly immune (7.57%) (Fig. 1A). Assignment of cell lineage identities were further validated by examining the expression of canonical marker genes associated with major cell types (Fig. 1B). Whereas expression patterns and levels varied for the canonical lineage markers, their expression was solely observed within clusters of their respective lineage. For example, EPCAM, SFTPB, and NXK2.1 are localized in the cluster classified as epithelium, whereas VWF, PECAM, and CLDN5 are expressed in the endothelial cluster (Fig. 1B).

**Figure 1.**
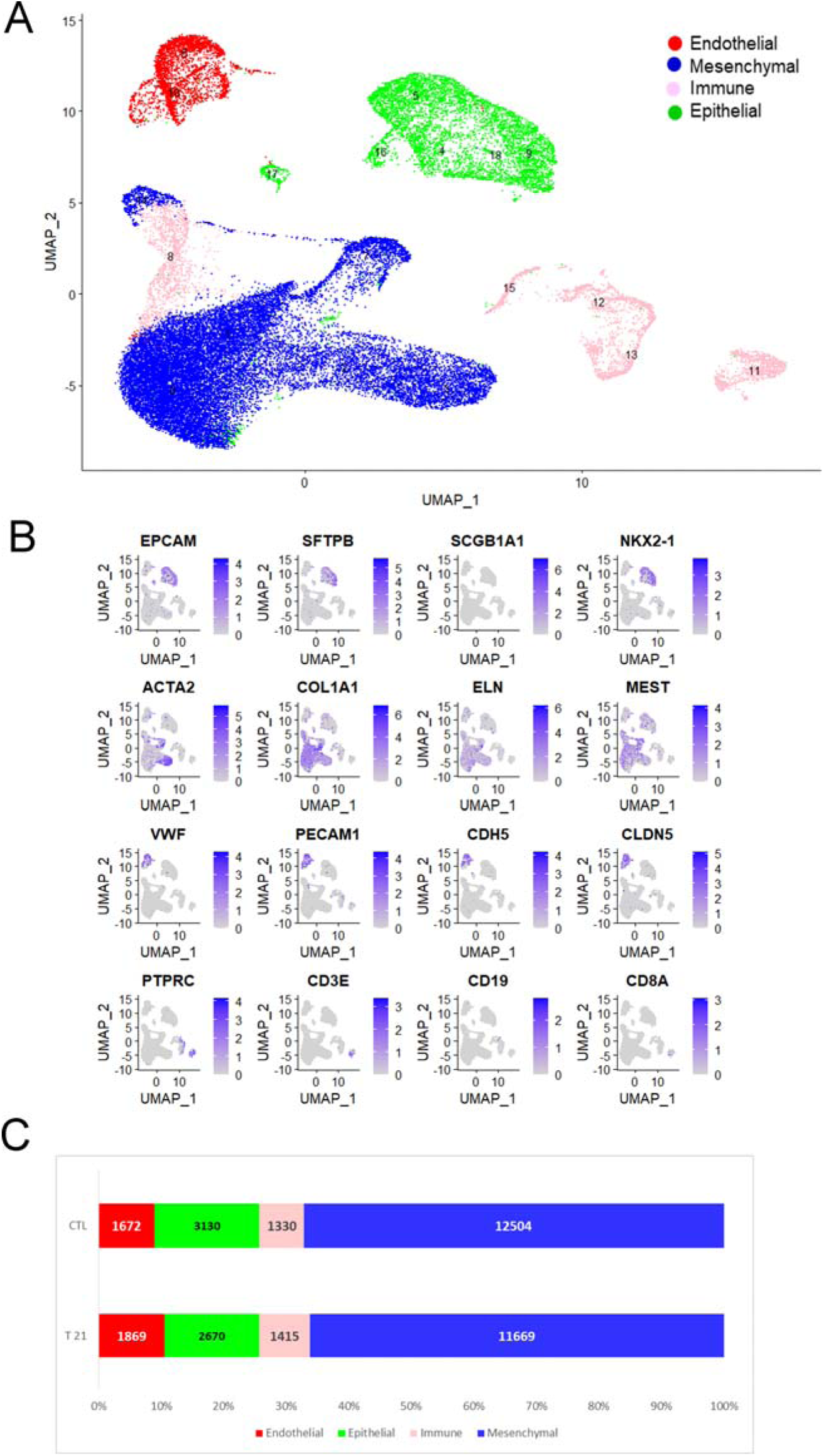
Single cell (sc) RNAseq of T21 and control non-T21 age matched lungs. Libraries from 9 samples, 4 non-T21 and 5 T21 lungs (all with anomalies) were sequenced, each within 17-19 weeks of gestation. A) Shown here are cells visualized using Uniform Manifold Approximation and Projection (UMAP). Each dot represents a single cell. Individual clusters are colored and annotated based on cell lineage association identified by ToppFun: Epithelium, Mesenchyme, Endothelium, and Immune. B) Cell lineages were validated using canonical marker genes. Expression patterns and level for individual canonical cell lineage markers are determined by unique molecular identifier (UMI) counts (depth of color). C) Cluster markers were organized into cell lineage type (Epithelial, Mesenchymal, Endothelial, and Immune) to demonstrate proportional composition in both non-T21 and T21 samples. No significant differences may be observed between the two groups.

Expression summary was assessed at both the diagnosis (T21 vs non-T21) and sample level, demonstrating no unique T21 clusters and no sample level bias (Fig. S1). Cell lineage proportionality was consistent when having separated and compared the T21 cells to the non-T21 (Fig. 1C). Although there are no significant or distinct cell populations observed in the developing T21 lungs as compared to the non-T21, we wanted to further assess proportional changes in the distribution of specific lung cell types and several differentially expressed genes that could account for the structural/developmental changes observed.

### Precocious epithelial cell differentiation in prenatal T21 lungs

We first focused on the epithelial lineage, given the most striking histopathological anomaly observed in prenatal T21 lungs is airway dilatation^2^. After sub-clustering the epithelial lineage, we identified a total of 12 distinct populations (Fig. 2A). For a complete list of epithelial cluster marker genes please see supplemental Table S4. The annotations represented all the major lung epithelial cell types: Basal Cells, Club Cells, Goblet Cells, Ciliated Cells, as well as the distal AT1 and AT2 populations. We acknowledge that many of these fully differentiated cell types do not yet exist in the mid-gestation fetal lung, but will refer to them using their annotated terms. We note that Cluster 11, which was annotated as Suprabasal-like, has many mesenchymal attributes. This could be indicative of a mixed lineage that is present during development, however further investigation outside the scope of this study is required. Although each cluster is unique, it was interesting to note that there was a redundancy amongst the cell type annotations (Fig. 2B). For example, both Clusters 2 and 5 are classified as AT1 cells whereas Cluster 6 is considered AT2 and Cluster 4 is considered an AT1/AT2 cell. The distinctiveness of each cluster was confirmed when looking at the expression level of specific canonical lung epithelial genes via dot and ridge plot, where we see that the expression of *HOPX*, *SFTPB* and *SFTPC* vary amongst the different clusters (Fig. 2B&C). Specifically, we note that AT2 (Cluster 6) differs from the AT1/AT2 (Cluster 4) population by having a higher average expression of *SFTPC*, whereas the AT1/AT2 cluster expresses *SFTPB* at a higher level. Furthermore, assessing the two distinct AT1 populations we note that there is markedly higher *HOPX* expression in AT1-(A) (Cluster 2) as compared to AT1-(B) (Cluster 5) but interestingly has a comparable expression level to that of the AT1/AT2 cluster (Cluster 4). Having established and annotated our unique epithelial populations, we note all subclusters are present for both T21 and non-T21, confirming there are no unique epithelial cell populations in the T21 lungs compared to non-T21 controls (Fig. S2A).

**Figure 2.**
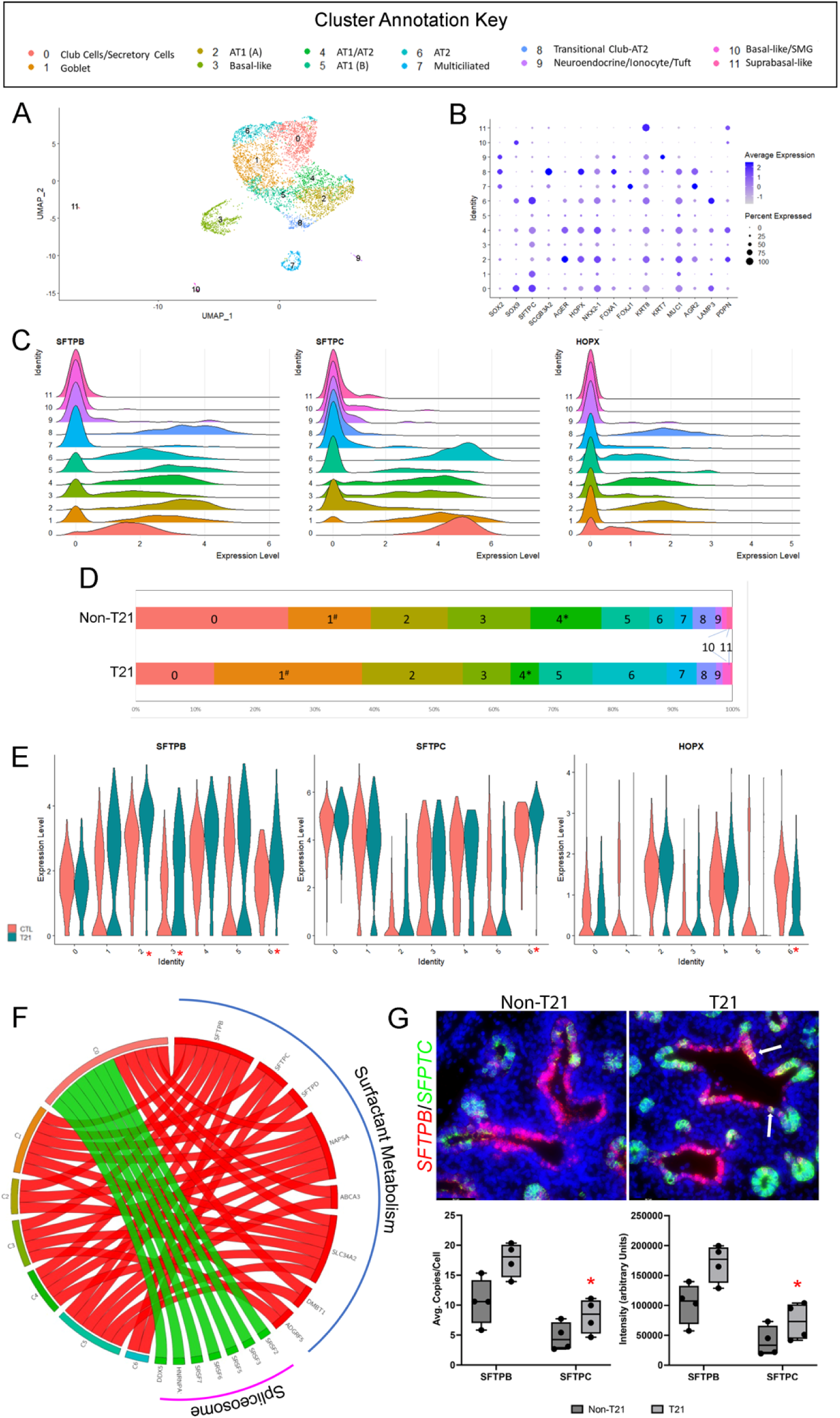
Trisomy 21 epithelial sub-clusters suggest accelerated differentiation as compared to controls. A) Uniform Manifold Approximation and Projection (UMAP) displays the 12 unique epithelial cell sub-clusters. Each dot represents a single cell and individual clusters are colored and annotated based on cell type associations identified by ToppFun, using marker genes for each individual cluster. B) Dot plot of cluster markers of canonical lung epithelial cell lineage validates these cells are of epithelial origin. *NKX2.1*, which is believed to be expressed in all developmental lung epithelial cells, is in almost each of the sub-clusters, whereas some more specialized markers, such as *FOXJ1*, known to be localized to ciliated cells, are in fact only found in the Multiciliated sub-cluster (Cluster 7). Within these populations the percentage of cells expressing the gene (using dot size) and the average expression level based on unique molecular identifier (UMI) counts (depth of color) were identified. C) Ridge plots of cluster markers characteristic of distal lung epithelium (*SFTPB, SFTPC, HOPX*) portray each cluster as unique. The x-axis represents the normalized expression level whereas the y-axis is indicative of the relative proportion of cells expressing the gene. D) Proportional differences in epithelial sub-clusters between non-T21 and T21 cohorts show T21 lungs have more Goblet cells (Cluster 1; p=0.07) whereas the non-T21 have significantly more “transitional” AT1/AT2 cells (cluster 4; p=0.01) (*p<0.05; #p<0.2). E) Violin plots of differentially expressed genes (*SFTPB, SFTPC, HOPX*) show an increase in several of the epithelial subclusters of T21 as compared to the non-T21. Significance (p<0.05) is indicated by * next to cluster number on x-axis. F) Pathway analysis comparing the T21 epithelial cells to the non-T21 shows upregulation (red) and downregulation (green) of several pathways. Surfactant Metabolism is shown to be upregulated in several of the T21 epithelial subclusters, suggestive of accelerated differentiation. Alternatively, other pathways, such as the Spliceosome Pathway, is downregulated in a single subcluster (Cluster 0). G) FISH of non-T21 control and T21 age and sex matched fetal lungs clearly demonstrate increased expression of both *SFTPB* and *SFTPC* within the epithelium of T21 lungs as compared to the non-T21, with several cells colocalized for both genes in the T21 lung (indicated by arrow; Fig. S1B). *SFTPC*: Copy Number p=0.0272; Intensity p=0.0347; *SFTPB*: Copy number p=0.1138, Intensity p=0.1089 (N=4 for each condition; Scale bar: 64.7um). Box Plots: center line, mean; box limits, upper and lower quartiles; whiskers, minimum to maximum

Although there are no unique clusters, when assessing the cellular composition of subpopulations between T21 and non-T21 lungs, there were clear proportional differences. Although not significant, T21 lungs showed a trend towards lower numbers of proximal progenitor cell types (Cluster 3, Basal; Cluster 0, Club/secretory), and towards more terminally differentiated proximal cell types (Cluster 1, Goblet; Cluster 7, Multiciliated). When considering distal epithelial cell types, T21 lungs showed an increased proportion of defined AT1 (Cluster 2) and AT2 (Cluster 6) cell populations, but a significant diminution in progenitor-like “transitional” AT1/AT2 cells (Cluster 4; p=0.01). These proportionality differences suggest precocious differentiation of epithelial cells occurs in T21 lungs^11^.

We next interrogated pathway activation in the epithelial cell clusters from developing T21 lungs. For each cluster, we identified differentially expressed genes and used those to assess dysregulated pathways. A complete list of differentially expressed genes for each epithelial cluster is presented in supplemental Table S5. Interestingly, we noted upregulation of the Surfactant Metabolism and downregulation of Spliceosome pathway (Fig. 2F). Due to the recognized importance of surfactant during lung development, we interrogated relevant genes of interest that were differentially expressed including *SFTPC, SFTPB, SFTPD*, and *ABCA3* (Fig. 2E, Fig. S2C). We noted that whereas *SFTPB* was significantly differentially upregulated in several of the T21 subclusters (Clusters 2, 3 and 6); *SFTPC* and *SFTPD* (Fig. S2C) were significantly differentially upregulated in unique subclusters: the AT2 cluster (cluster 6) and AT1 (A) (cluster 2) respectively. Furthermore, our data showed significantly increased expression of *ABCA3* in the “AT2” cluster, a gene recently implicated as a marker of mature AT2 cells^12, 13^ (Fig. S2C). Taken together the increased differential expression of several surfactant proteins, as well as the increase of ABCA3 in the AT2 cluster strongly suggests premature differentiation and/or maturation of distal epithelial cells in T21 lungs.

These results were validated using combinatorial FISH-IF to quantify the expression of the different genes in context to their localization. Specifically, we note higher expression of *SFTPC* within the distal epithelium of the T21 lungs as compared to the non-T21 sex and age matched controls and stronger presence of *SFTPB* in the proximal epithelium, both which were determined by the quantification of average transcription copy number per cell as well as expression intensity (*SFTPC*: Copy Number p=0.0272; Intensity p=0.0347; *SFTPB*: Copy number p=0.1138, Intensity p=0.1089). Of interest is that the T21 lungs have a larger number of double positive *SFTPC/SFTPB* cells within the proximal epithelium (Fig. 2G), which was suggested in the single cell data (p=0.1532; Fig. S2B). Furthermore, although the percentage of epithelial cells positive for *ABCA3* is comparable between T21 and non-T21 lungs, there appears to be on average more transcripts per cell (p=0.1310) in the T21 lungs with a stronger intensity (p=0.0669), validating that *ABCA3* expression is in fact greater within the epithelium of developing T21 lungs (Fig. S2C).

### Dysregulated mesenchymal ECM production is associated with compromised branching

Proper lung branching morphogenesis is a dynamic process that requires cross-talk between the epithelium and mesenchyme, and can be driven by spatiotemporal deposition of extracellular matrix. Therefore, to better understand the branching defects observed in the developing T21 lung, we next investigated the mesenchymal compartment. We determined that the fetal human lung has 10 unique mesenchymal subclusters (Fig. 3A). A complete list of mesenchymal cluster marker genes can be found in supplemental Table S6. There was one non-specific mesenchymal cluster, Cluster 1, which expressed markers of all major cell lineages (including Epithelial, Endothelial, and Immune), suggesting the presence of possibly several “multipotent” cell populations in the developing lung that remain poorly described.

**Figure 3.**
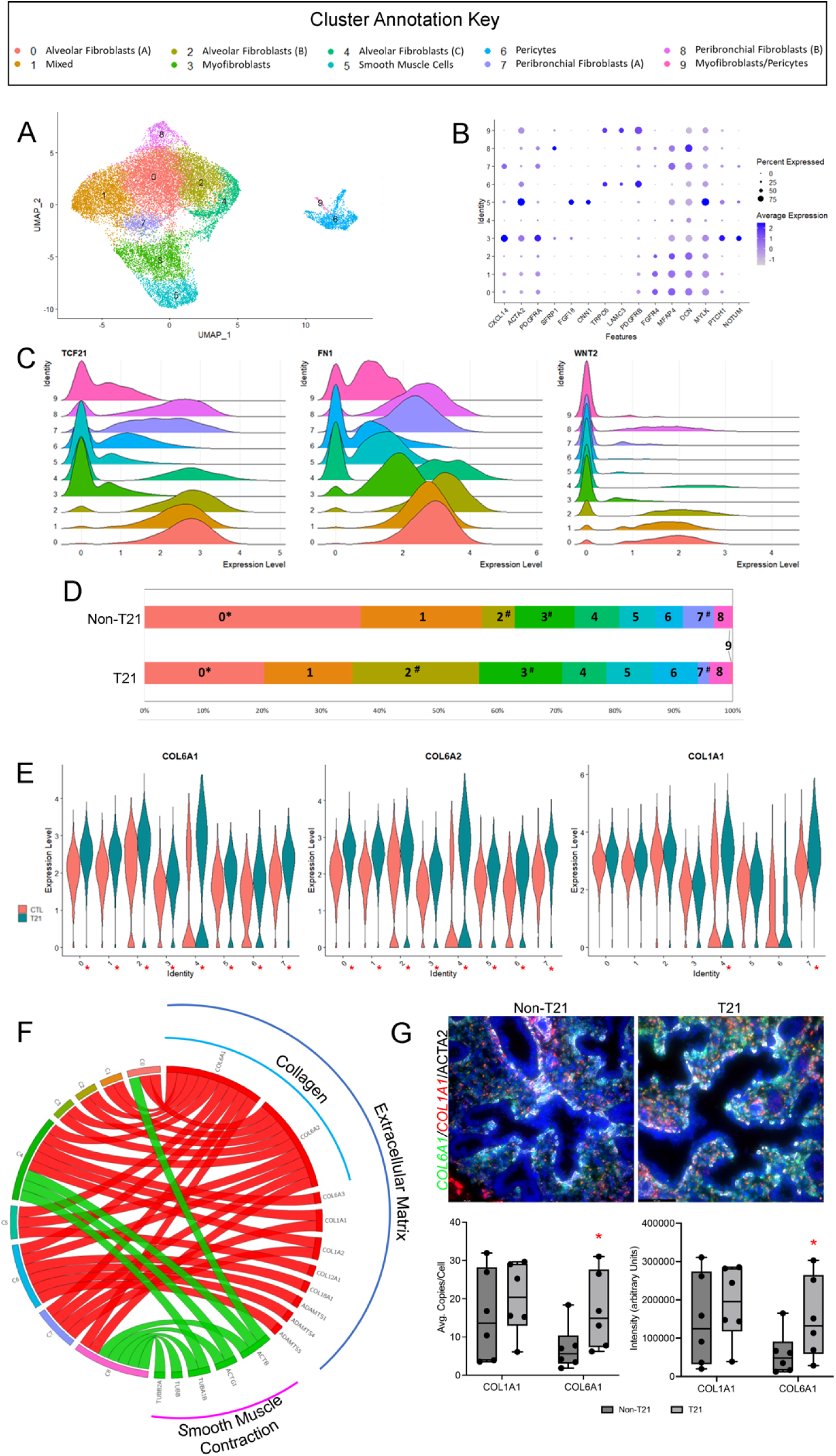
Dysregulated ECM contributes to compromised T21 branching. A) Uniform Manifold Approximation and Projection (UMAP) displays the 10 unique mesenchymal cell sub-clusters. Each dot represents a single cell and individual clusters are colored and annotated based on cell type associations identified by ToppFun, using marker genes for each individual cluster. B) Dot plot of cluster markers of canonical lung mesenchymal cell lineage validates these cells are of mesenchymal origin. *DCN*, which is a universal developmental lung mesenchymal cell, is in almost each of the sub-clusters, whereas some more specialized markers, such as *PDGFRB*, which is a known marker of pericytes, are primarily found in the Pericyte sub-cluster (Cluster 6). Within these populations the percentage of cells expressing the gene (using dot size) and the average expression level based on unique molecular identifier (UMI) counts (depth of color) were identified. C) Ridge plots of cluster markers that are strongly associated with the lung mesenchyme (*TCF21, FN1, WNT2*) clearly portray each cluster as unique, regardless of redundancies in cell annotations. The x-axis represents the normalized expression level whereas the y-axis is indicative of the relative proportion of cells expressing the gene. D) Proportional differences in mesenchymal sub-clusters between non-T21 and T21 cohorts show T21 lungs have a reduced population of Peribronchial Fibroblasts (Cluster 7; p=0.110). They also have a trend towards a larger population of Cluster 2 Alveolar Fibroblasts (B) (p=0.162). Conversely, non- T21 lungs have a significantly larger population of Cluster 0 Alveolar Fibroblasts (A) (p=0.022) (*p<0.05; #p<0.2). E) Violin plots of differentially expressed genes (*COL6A1, COL6A2, COL1A1*) show an increase in several of the mesenchymal subclusters of T21 as compared to the non-T21. Significance (p<0.05) is indicated by * next to cluster number on x-axis. F) Pathway analysis comparing the T21 mesenchymal cells to the non- T21 shows upregulation (red) and downregulation (green) of several pathways. Both the Collagen Pathway and the Extracellular Matrix pathway are upregulated in several of the T21 mesenchymal subclusters, with overlap between the two pathways. Alternatively, the Smooth Muscle Contraction pathway is shown to be downregulated in the Peribronchial Fibroblast (B) cluster (Cluster 8). G) FISH of non-T21 control and T21 age and sex matched fetal lungs clearly demonstrate increased expression of both *COL6A1* and *COL1A1* within the mesenchyme of T21 tissues as compared to the non-T21, with several cells colocalized for both genes in the T21 (Fig. S3B). *COL6A1*: Copy number p=0.0168, Intensity p=0.0190; *COL1A1*: Copy number p=0.2866, Intensity p=0.2979 (N=6 for each condition; Scale bar: 64.7um). Box Plots: center line, mean; box limits, upper and lower quartiles; whiskers, minimum to maximum

Some mesenchymal clusters display redundant cell type annotations but are clearly unique as determined by the expression of several canonical lung mesenchymal genes (Fig. 3B). While alveoli do not exist at this stage of human fetal lung development, Clusters 0, 2 and 4 are all classified as Alveolar Fibroblasts. Each of these clusters, however, have differing levels of expression of both *FGFR4* and *FN1* (Fig. 3B&C). Clusters 7 and 8 are both classified as Peribronchial Fibroblasts, however express differing levels of several genes, including *CXCL14, SFRP1, PDGFRB*, and *WNT2*. Finally, whereas Cluster 3 is solely labeled as Myofibroblasts and Cluster 6 as Pericytes, Cluster 9 is labeled as Myofibroblasts/Pericytes. When comparing Cluster 3 to 9 there are differences in gene expression for *PDGFRa, MFAP4, PTCH1*, and *FN1;* whereas comparing Cluster 6 to 9 demonstrates differing gene expression of *LAMC3, PDGFRb*, and *TCF21*.

We confirmed that no unique mesenchymal cell populations existed in the T21 lungs (Fig. S3A). Proportional differences for some mesenchymal subpopulations were observed and were consistent with our prior observations indicating reduced branching in T21 fetal lungs^3, 9^. We note a reduced population of Peribronchial Fibroblasts (Cluster 7; p=0.110), which are associated with airways. Non-T21 lungs have a significantly larger population of Cluster 0 Alveolar Fibroblasts (A), whose cluster markers are indicative of a naive fibroblast (p=0.022). Conversely, T21 lungs have a trend towards a larger population (p=0.162) of Cluster 2 Alveolar Fibroblasts (B), which are marked by expression of several collagen genes shown to inhibit lung branching morphogenesis (i.e *COL6A1, COL6A3, COL1A1*, etc.)^14^. This suggests there is a phenotypic shift in T21 lung mesenchymal cells from a less differentiated fibroblast to a more differentiated, branching-constraining fibroblast (Fig. 3D).

We next interrogated differentially expressed genes in each of the 10 mesenchymal subclusters. For a complete list of differentially expressed genes for each mesenchymal cluster please see supplemental Table S7. We observed an upregulation of several interstitial matrix associated fibrillar collagens, whose role is to contribute to the mechanical strength and framework of the tissue. *COL6A1* and *COL6A2* were consistently upregulated in the mesenchymal subclusters (Fig. 3E). Pathway analysis of differentially expressed genes confirmed upregulation of ECM and Collagen pathways in several mesenchymal subclusters. Interestingly, this analysis also identified downregulation of the Smooth Muscle Contraction pathway (Fig. 3F; Fig. S3D), which is primarily observed in the Peribronchial Fibroblasts (B) (Cluster 8).

We quantified the expression of different ECM genes in context to their localization using combinatorial FISH-IF. *COL6A1* and *COL6A2*, located on HSA21, were significantly increased in both the copy number per cell as well as signal intensity in T21 lungs as compared to controls (*COL6A1*: Copy number p=0.0168, Intensity p=0.0190; *COL6A2*: Copy number p=0.0346, Intensity p=0.0322) (Fig. 3G; Fig. S3C). Conversely, there was no significant increase in either copy number or signal intensity for *COL1A1*, but we did observe a trend of an increase in both copy number (p=0.2866) and Intensity (p=0.2979). This may be due to *COL1A1* being upregulated only in Clusters 4-Alveolar Fibroblasts (C) and 7-Peribronchial Fibroblasts (A). Finally, we note there was a significantly larger population of double positive *COL6A1/COL1A1* expressing cells in the T21 lungs as compared to the non-T21 lungs (p=0.0289; Fig. S3B).

### Altered T21 lung vascular development is associated with aberrant type I interferon (IFN) signaling in endothelial cells

We and others have demonstrated that the T21 lung often presents with abnormal vascular growth in utero^9, 15^. Whereas an upregulation of certain anti-angiogenic factors has been reported in these lungs, there is limited information describing the ontogeny of these defects. Focusing specifically on the endothelial cell population, we determined a total of 11 distinct endothelial sub-populations (Fig. 4A). A complete list of endothelial cluster marker genes is shown in supplemental Table S8. Much like the other cell lineages, the annotations represented all the major lung endothelial cell types, including the newly described aCaps (Cluster 2). However, there were two mixed populations (Cluster 3&4), as well as clusters classified as non-endothelial (Cluster 8&10), again revealing the cellular diversity observed in the developing human lung. When assessing the expression of canonical endothelial markers, such as *PECAM1* and *CLDN5*, each of the sub-clusters was positive, thus establishing their endothelial nature (Fig. 4B).

**Figure 4.**
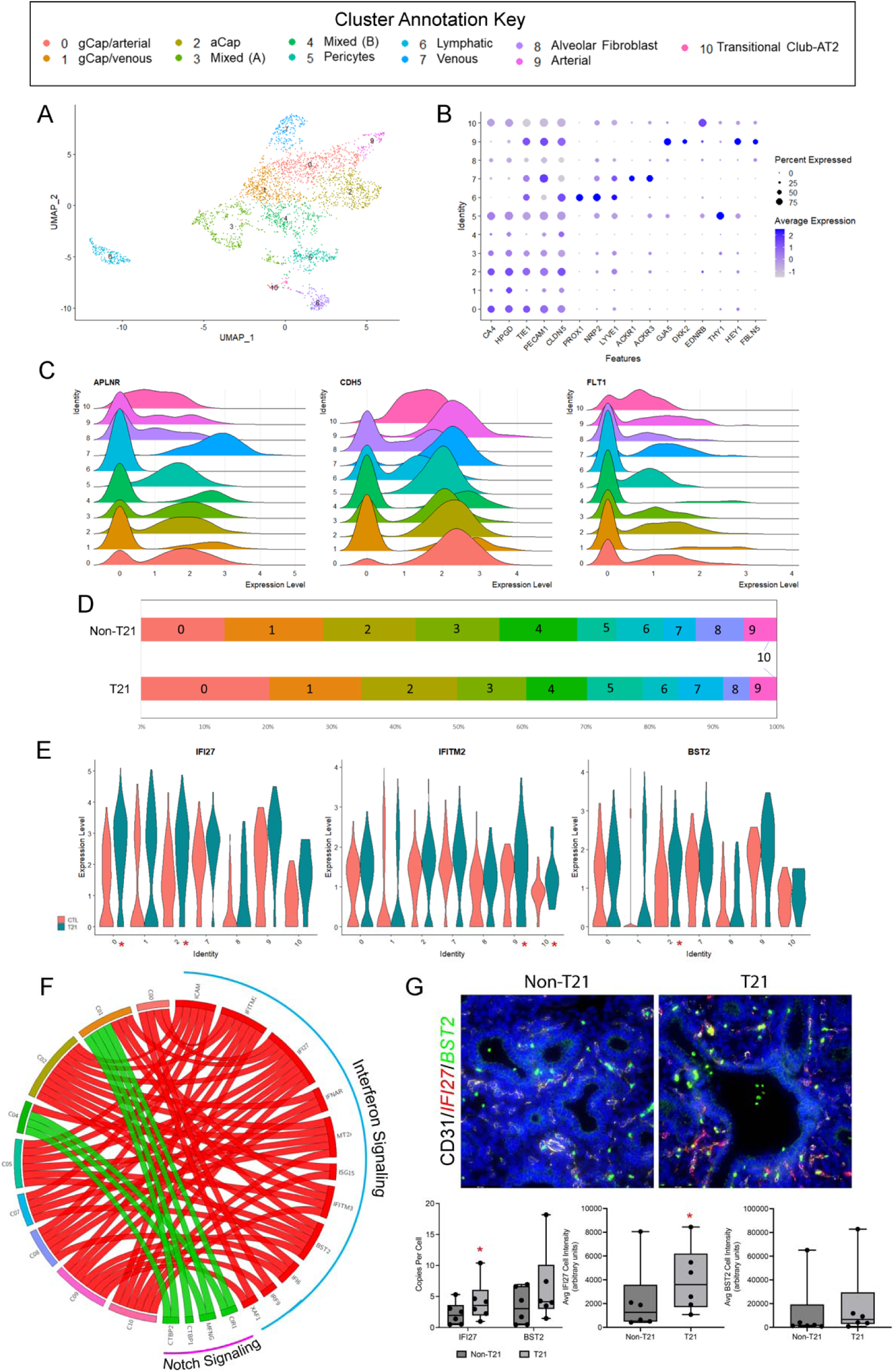
Upregulated Type I IFN signaling disrupts endothelium in prenatal T21 lungs. A) Uniform Manifold Approximation and Projection (UMAP) displays the 11 unique endothelial cell sub-clusters. Each dot represents a single cell and individual clusters are colored and annotated based on cell type associations identified by ToppFun, using marker genes for each individual cluster. B) Dot plot of cluster markers of canonical lung endothelial cell lineage validates these cells are of endothelial origin. *PECAM1* and *CLDN5*, universal developmental lung endothelial cell markers, are expressed in each of the sub-clusters, whereas some more specialized markers, such as *PROX1*, a known lymphatic marker, are solely found in the Lymphatic sub-cluster (Cluster 6). Within these populations the percentage of cells expressing the gene (using dot size) and the average expression level based on unique molecular identifier (UMI) counts (depth of color) are identified. C) Ridge plots of cluster markers that are strongly associated with the lung endothelium (*APLNR, CDH5, FLT1*) clearly portray each cluster is unique, provided their differing levels of expression amongst the clusters. The x-axis represents the normalized expression level whereas the y-axis is indicative of the relative proportion of cells expressing the gene. D) Interestingly, unlike that observed in the epithelial and mesenchymal cells, proportional differences in endothelial sub-clusters between non-T21 and T21 cohorts show that none of the clusters are significantly different. E) Violin plots of differentially expressed genes (*IFI27, IFITM2, BST2*) show an increase in several of the endothelial subclusters of T21 as compared to the non-T21. Each of these are downstream targets in the type I IFN signaling pathway. Significance (p<0.05) is indicated by * next to cluster number on x-axis. F) Pathway analysis comparing the T21 endothelial cells to the non-T21 shows upregulation (red) and downregulation (green) of several pathways. The Interferon Signaling pathway is upregulated in several of the T21 endothelial subclusters. Alternatively, the Notch Signaling pathway is shown to be downregulated primarily in the gCap/Venous cluster (Cluster 1). G) FISH of non-T21 control and T21 age and sex matched fetal lungs clearly demonstrate increased expression of both type I IFN markers *IFI27* (red) and *BST2* (Green) within the endothelium (CD31-White) of T21 lung as compared to the non-T21, with several cells colocalized for both genes in the T21 (Fig. 3SB). *IFI27*: Copy number p=0.0470, Intensity p=0.0163; *BST2*: Copy number p=0.1471, Intensity p=0.0777 (N=6 for each condition; Scale bar: 64.7um). Box Plots: center line, mean; box limits, upper and lower quartiles; whiskers, minimum to maximum

Unlike the replicated annotations observed in both the epithelium and mesenchyme, each of the endothelial clusters were uniquely annotated. There was only a slight redundancy between Clusters 0 and 1, which were both co-classified, with gCap being the primary annotation. However, when assessing the level of expression of certain well-established endothelial markers, it is evident that they are quite unique populations (Fig. 4C). Specifically, Cluster 0 has more *APLNR* and *CDH5* expressing cells than Cluster 1, but Cluster 1 appears to have cells that express both genes at a higher level (Fig. 4C). Additionally, there is partial redundancy between Clusters 0 and 9, which both have an arterial association, as well as between Clusters 1 and 7, which both have a venous association. However, the ridge plots clearly demonstrate that these populations are in fact unique compared to one another as there is greater expression of *FLT1, APLNR* and *CDH5* in Cluster 7 as compared to 1, and greater expression of *APLNR* in cluster 0 as compared to 9. We confirmed that no unique endothelial cell populations exist in the T21 lungs (Fig. S4A). Unlike in the Epithelial and Mesenchymal lineages, it was interesting to note that there are no significant proportional differences in any of the endothelial sub-types (Fig. 4D).

Using differentially expressed genes between the two cohorts, we performed pathway analysis to assess endothelial cell differences in the developing T21 lungs that are associated with the vascular abnormalities we have described ^9, 15^. For a complete list of differentially expressed genes for each endothelial cluster please see supplemental Table S9. Interestingly, we saw upregulation of the type I IFN Signaling Pathway and downregulation of the Notch Signaling pathway (Fig. 4E&F). Specifically, we noted several IFN related genes that were significantly differentially expressed in several of the sub-clusters including: *IFI27, BST2, IFITM2*, and *IFITM3* (Fig. 4E; Fig. S4C). This upregulation is of interest given that high type I IFN inhibits VEGF induced development of capillary like structures^16, 17^ and is associated with a decrease of endothelial progenitor cells^18^. This, in conjunction with downregulated Notch Signaling, a well-established pathway involved in the cell fate, differentiation, and angiogenesis of vascular endothelial cells, are possibly implicated in the vascular “simplification” phenotype observed in the T21 lung^9, 15^.

Again, we validated these results using combinatorial FISH-IF to quantify the expression of different type I IFN associated genes in context to their localization. In both the non-T21 and T21 lungs *BST2* and *IFI27* are primarily localized in the endothelium. Comparing the expression levels between the two cohorts, we note that the average transcript number and intensity per endothelial cell of *IFI27* is significantly greater in T21 as compared to non-T21 (*IFI27*: Copy Number: p=0.0470; Intensity: p=0.0163). Additionally, there is an upwards trend of *BST2* transcript number and intensity in the T21 lungs as compared to the non-T21 (*BST2*: Copy Number: p=0.1471; Intensity: p=0.0777). However, when looking at the co-expression of these two genes, we see a significantly larger population of double positive *IFI27/BST2* expressing endothelial cells in the T21 lungs as compared to the non-T21 (p=0.0230; Fig. S4B).

## Discussion

Adult and pediatric respiratory morbidity has long been recognized as a complication of Down Syndrome. The gross structure of the mature T21 lung is quite comparable to that of a non-T21 lung, but may present with mild alveolar, airway and vascular changes. We recently described a fetal origin for lung histopathological abnormalities in T21^9^. The goal of the studies reported here was to comprehensively catalog cellular and molecular alterations in the fetal T21 lung, coincident with the onset of histological changes. It comes as no surprise that there is a similar composition of cell types at the lineage level (epithelial, mesenchymal, endothelial, immune) in T21 and non-T21 lungs (Fig. 1). However, it is interesting to note in our data presented herein, that multiple differences in cellular proportions are observed dependent upon individual lineages. Furthermore, unique differences in gene expression are also observed in each lineage. Our data uncover an early and complex effect of triplication of HSA21 upon lung morphogenesis and provide a foundational description of molecular changes at the cellular level.

In the epithelial lineage we noted strong evidence of precocious cellular differentiation in the T21 lungs as compared to the non-T21 controls^11^. These observations are of interest given that DS has been described as a progeroid syndrome, characterized by accelerated maturation or aging^19–21^. Individuals with DS start aging prematurely, and present with conditions more characteristic of the geriatric population^21, 22^. These signs of early aging have been described to affect primarily the neurological system, with high prevalence of neurodegenerative disorders (i.e. dementia and Alzheimer’s disease) as well as the immune system^23^. It has been demonstrated that this accelerated aging may be associated to genome wide perturbation in epigenetic regulation^24, 25^. When conducting pathway analysis on our 12 unique epithelial populations (epithelial sub-clusters) we determined an upregulation of surfactant metabolism in the T21 lungs, which was confirmed by increased differential expression of several surfactant proteins (*SFTPB, SFTPC, SFTPD*) (Fig. 2E; Fig. S2C). Furthermore, our single cell data showed significantly increased expression of *ABCA3*, a gene recently shown to be a marker of mature AT2 cells^12^, in two clusters. ABCA3 is expressed in the human fetal lung normally after 28 weeks gestation^12, 13^; however, we observe a robust expression at 18 weeks gestation in T21 in the AT2 Cluster (Cluster 6) (Fig. S2C). This concept of premature maturation may further be extrapolated from our previous findings where we demonstrated significantly decreased proliferation in the human fetal T21 lungs as compared to age and sex matched non-T21 lungs in conjunction with decreased proximal airway SOX2 expression and abnormal airway smooth muscle cell (ASMC) patterning^9^. Altogether these data support that premature differentiation and precocious development is occurring in the epithelium of T21 fetal lungs.

Further evidence of accelerated differentiation is also seen in the mesenchymal lineage. The two populations that were proportionally different in the mesenchymal cells were both annotated as Alveolar Fibroblasts, with Cluster 0 being larger in the non-T21 lungs and Cluster 2 being larger in the T21 lungs. When assessing their cluster markers, it appears that Cluster 0 is a less differentiated fibroblast as compared to the more differentiated collagen producing fibroblasts of Cluster 2. This advanced differentiation of the mesenchymal cells could ultimately influence the compromised branching previously described in the prenatal T21 lung^9^. The significant upregulation of both *COL6A1* and *COL6A2* in the T21 lungs is of particular interest given that deletion of *COL6* has been shown to limit airway branching^14^. Additionally, we noted a smaller population of Peribronchial Fibroblasts (Cluster 7) in the T21 lungs, as well as downregulation of the smooth muscle contraction pathway in a second Peribronchial Fibroblast population (Cluster 8). It has been previously demonstrated that smooth muscle cells play an important role in branching during fetal lung development^26^. Furthermore, T21 fetal lungs are known to present with a thinner and interrupted layer of smooth muscle cells around the dilated airways^9^. Thus, taken together, the upregulation of both the Collagen and ECM pathways in T21 lungs, as well as the decreased Peribronchial fibroblasts and smooth muscle cell contraction, suggest that dysregulated ECM components of the fetal T21 is likely contributing to the compromised branching observed.

Whereas the ECM and collagen pathways are expected to be affiliated with the mesenchymal cell population, it was somewhat surprising to note that the IFN signaling pathway was most prominent in the endothelial lineage. The type I IFN pathway is one of the major signaling cascades consistently activated in T21^10^. This was further confirmed in T21 fetal lungs, where bulk sequencing demonstrated an appreciable upregulation of IFN pathway genes^9^. With single cell sequencing, we have now demonstrated that this aberrant IFN signaling is primarily localized in the endothelium in the fetal T21 lung (Fig. 4F&4G). Although there are no proportional differences in any of the endothelial clusters (Fig. 4D), each cluster differentially expressed several downstream type I IFN targets: *IFI27, IFITM2, BST2*, and *IFITM3* (Fig. 4E). While multiple groups have reported congested capillaries, lymphatic dilatation, and muscularized arteries in the developing T21 lung^9, 27^, the nature of these defects and the different endothelial cell populations contributing to such defects are yet to be understood. Upregulation of type I IFN signaling has been previously shown to inhibit endothelial cell proliferation and block angiogenesis^16^. Our single cell data demonstrated a downregulation of the Notch signaling pathway in endothelial cells, which is known to be a key regulator for multiple steps in angiogenesis/vasculogenesis, from sprouting to specification^28^. Ultimately, the combined upregulation of type I IFN activity and the associated dysregulation of Notch signaling are likely contributing to the observed compromised vasculature. Interestingly, altered vascular development has been shown to also affect fetal lung branching^29^.

As we have previously demonstrated, and validated herein, the prenatal human lung presents with several immune cell populations^7^. It has been well-established that individuals with DS have numerous immune defects^30^. Therefore, it is interesting to note that in our dataset the immune cell population showed the least prominent differences between the T21 and non-T21 lungs. This may be due to the mid-gestation lung being a poor model for the immune cells, given their sparsity. However, we do note that there are a total of 12 unique immune cell populations that are detected in both cohorts (Fig. S5). For a complete list of immune cell cluster marker genes please see supplemental Table S10. Whereas there were differentially expressed genes between T21 and non-T21 within each cluster when assessed via unadjusted p-values, only a small number were differentially expressed with an adjusted p-value. Therefore, we chose to focus our current analysis on the resident cell populations. An expanded assessment of immune cells in the postnatal T21 lung will be necessary to draw definitive conclusions.

In this study we present a comprehensive cellular atlas of the T21 lung. Specifically, we describe changes in gene regulatory networks at the single cell level in T21 fetal lungs presenting with developmental anomalies. We demonstrate that there are certain cell populations that differ significantly in proportionality and many others displaying unique patterns of differential gene expression. These data provide critical insight as to how specific pathways and cellular processes are being altered in different cellular lineages, and how these may lead to advanced cellular differentiation and altered lung development in the T21 lung. This benchmark cellular atlas of T21 lungs should greatly facilitate efforts to ameliorate respiratory morbidity in patients with Down Syndrome.

## Online Methods

### Human Tissue

The human fetal lung tissues used in this study were collected under IRB approval (USC-HS-13-0399 and CHLA-14-2211) provided by the USC fetal tissue biobank and the University of Washington Birth Defects Research Laboratory. Following informed consent, de-identified human fetal samples were collected. The only information provided was gestational age and whether there were any known genetic conditions [chromosomal trisomy 21 (T21), T18, T13] or gestational complications such as low amniotic fluid. T21 diagnosis was made at the different collection centers by routine prenatal diagnostic methods. Lungs were received as fresh whole lobes and processed either immediately for single cell suspension or fixed in 4% PFA for immunohistochemical analyses. Histologic review of T21 lungs and age-matched controls was carried out by an experienced pediatric pulmonary pathologist (GD). Histopathological analyses of T21 lungs as compared to age matched control lungs are detailed in supplemental Table S1.

### Single cell suspension preparation

Human fetal lung tissues were dissociated using the Miltenyi Neural Tissue Dissociation Papain-based Kit (Cat.130-092-628) according to the manufacturer’s instructions alongside additional adaptations. Prior to tissue dissociation, all plasticware to be used throughout the protocol (i.e falcon tubes, pipette tips, cell strainer etc.) were rinsed/coated with filtered 1%BSA in HBSS. From each lung, 400 mg of tissue was weighed out and placed on a clean glass petri dish containing chilled HBSS and minced (1-2 mm sized pieces) using a sterile razor blade. Minced lung with HBSS was then transferred to a 50ml Falcon tube and spun at 500G for 5 minutes. Excess HBSS was then decanted, and lung pieces were resuspended in Neural Tissue enzyme Mix 1 (50uL Enzyme P + 1900uL Buffer X) and transferred to a gentleMACs C tube (Cat: 130-093-237; gentleMACs dissociator compatible tubes). Lung was then further dissociated with the Miltenyi gentleMACs dissociator (Cat: 130-093-235) using the manufacturer-set dissociator program “mouse tumor implant program-01.01”. Sample was then incubated in 37°C cell incubator, with cap slightly loosened for 15 minutes (for all subsequent incubation steps note that the cap is slightly loosened). Enzyme Mix 2 (10uL Enzyme A and 20uL Buffer Y) was then added to the tissue and manually agitated, first with a P1000 pipette tip with the tip cut followed by manual agitation with an uncut P1000 pipette tip. Sample was then placed back into 37°C cell incubator for 10 minutes. After incubation, sample was fluxed with P1000 tip followed by a P200 tip, then placed back in incubator for an additional 10 minutes. Sample was again fluxed with a P200 and placed in incubator for an additional 10 minutes (these two steps were repeated for an additional 3-4 times until achieve a single cell suspension). During these incubation steps a 70um cell strainer was coated with 1% BSA and rinsed with HBSS. Cell solution was then passed through prepared 70um cell strainer. C tube was then rinsed with 3mL HBSS and passed through strainer, and strainer was rinsed further with an additional 2mL HBSS. Strained cell suspension was then centrifuged at 500G for 5 minutes followed by decanting. Cell pellet was then incubated in Red Blood Cell Lysis Buffer (3 minutes at room temperature), diluted with HBSS (4 HBSS:1 RBC), and centrifuged at 500G for 5 minutes followed by decanting. Cell pellet was subsequently resuspended in 2mL 1% BSA and centrifuged at 500G for 5 minutes followed by decanting (this step was repeated a second time). Cells were then resuspended in 1%FBS DMEM/F12 at a concentration of 1000 cells/μL for single cell RNA library preparation with all subsequent cells frozen down in 90% FBS/10% DMSO solution.

### Single Cell Sequencing

Single cell capture and library production was performed on the Chromium 10X Genomics system with version 3 chemistry according to the manufacturer’s instructions. Library quality control was performed on the Agilent Bioanalyzer 2100 using the High Sensitivity DNA Kit (Agilent Technologies, cat. no. 5067-4626). Sequencing was performed on an Illumina NovaSeq, with read alignment to GRCh38. All single cell sequencing data analysis was performed using Seurat v3.2. Filtered data were log transformed, scaled, integrated using reverse Principal Component Analysis (rPCA), clustered and represented by Uniform Manifold Approximation and Projection (UMAP) dimensionality reduction. Differential expression was defined using a non-parametric Wilcoxon rank sum test at a significance level of p<0.05 corrected for multiple testing. Pathway analysis and cell type association was performed using ToppGene Functional Annotation tool (ToppFun).

### Fluorescent in situ hybridization (FISH)/Immunofluorescent Staining

Lung tissues (T21 and non-T21 age and sex matched controls) were fixed in 4% PFA for 24-48 hours, gradually dehydrated in a series of ethanol washes, followed by tissue clearing in xylene. Tissues were then paraffin embedded, sectioned at 5um, and dried overnight at 37°C. Sections were subsequently baked at 60°C for 1 hour in preparation for combinatorial ACD RNAScope In Situ Hybridization (FISH) and immunofluorescent (IF) staining, which was performed as previously described^7^.

Briefly, tissues were de-paraffinized in a series of xylene and ethanol. Ready-to-use ACD hydrogen peroxide (Ref:322335) was added to the tissue and kept at room temperature (RT) for 10 minutes. Slides were then submerged in boiling ACD target retrieval solution (Ref:322000) for 15 minutes and further incubated with ACD Protease plus solution (Ref:322331) at 40°C for 25 minutes. Sections were then incubated with assigned probes (supplemental Table S2) at 40°C for 2 hours (Channel 2/3 probes were diluted into a Channel 1 probe at a ratio of 1:50). Slides were then submerged in 5X SSC Buffer (Ref: BP1325) at RT overnight. The following day signal amplification and detection was developed using the ready-to-use reagents within the ACD RNAScope Multiplex Fluorescent V2 Assay (Cat. No. 323100) per the manufacturer’s instructions with OPAL Fluorescent Dyes used for labeling (Supplemental Table 3). Upon completion of FISH, slides were washed for 3 minutes with 1xTBST, blocked in 3% bovine serum album/5% Normal Goat Sera/0.1% Triton-X100 in TBS Blocking Solution (1 hour at RT) and incubated with primary antibodies at 4°C overnight (supplemental Table S3). The following day, slides were washed twice for 5 minutes in 1xTBST, and slides were then incubated with species appropriate fluorescent conjugated secondary antibodies (supplemental Table S3) for 1 hour at RT. Slides were then washed twice for 5 minutes in 1x TBST, counterstained with DAPI (Invitrogen Ref: D3571) and mounted with Prolong Diamond Antifade Mounting Media (Invitrogen Ref: P36961).

### Image Analysis

Images were obtained using a Lecia THUNDER inverted microscope. The different combinations of combinatorial FISH/IF were performed on up to 6 different lungs for each condition (T21 and non-T21). Sections were imaged at 40x magnification, with 10 images were assessed per sample. Using HALO Image Analysis Platform Version 3.5.3577 (Indica Labs,Inc) each pair was individually assessed to assure accurate quantification of RNA copy number, intensity, and protein expression.

### Statistical analyses for FISH Quantification

Due to the limited number of human samples, non-parametric Mann–Whitney and Kruskall–Wallis tests were used as appropriate. A paired Student’s t-test was used to detect significant differences between non-T21 and T21 tissues that were age and sex matched. The results were considered significant if p ≤ 0.05.

## Author Contributions

SB, TJM, DAA, and SD designed the research studies; SB, CC, and SD conducted the experiments; SB, CC, GD and SD acquired data; SB, TJM, DAA, and SD analyzed the data; GD and IG provided human tissues and input on data; SB, CC, GD, IG, TJM, DAA, and SD wrote edited and approved the final manuscript. No honorarium or other form of payment was given to anyone to produce the manuscript.

## Data Availability

The datasets generated during and/or analyzed during the current study are available in the NCBI SRA/GEO repository (Accession Number Pending).

## Acknowledgements

We thank Melissa L. Wilson (Department of Preventive Medicine, University of Southern California) and Family Planning Associates for coordinating fetal tissue collection. We also thank Dr. Brendan H Grubbs and Matthew E Thornton (Department of Obstetrics and Gynecology, Maternal Fetal Medicine Division, Keck School of Medicine, University of Southern California) and the Birth Defects Research Lab (BDRL) at Washington: Laboratory of Developmental Biology was supported by NIH Award Number R24HD000836 from the Eunice Kennedy Shriver National Institute of Child Health & human Development. We acknowledge the University of Rochester Genomics Research Center (GRC) for completing the Single Cell RNA sequencing (sc-RNA-Seq).

