## Supplemental Data for "A Trisomy 21 Lung Cell Atlas"

Supplemental Tables/Figures:

Table S1. Histopathological Assessment of Samples Used

| Samples | Gestational Age | Sex | Genetic Anomalies | Dilated DB and Acinar Tubules | How used |
|---|---|---|---|---|---|
| 3416 | 106D | M | | 0 | Single Cell |
| 3442 | 106D | F | T21 | 1+ | Single Cell |
| 3463 | 108D | F | | 0 | Single Cell |
| 3675 | 108D | F | | 0 | Single Cell |
| H28433 | 122D | F | T21 | 2+ | Single Cell |
| H28758 | 108D | M | | 0 | FISH/IF |
| H28790 | 113D | M | T21 | 2+ | FISH/IF |
| H28670 | 108D | M | | 0 | Fish/IF |
| H28406 | 108D | M | T21 | 1+ | Single Cell FISH/IF |
| 69 | 113D | F | | 0 | FISH/IF |
| 3412 | 113D | F | T21 | 1+ | Single Cell FISH/IF |
| H28554 | 120D | F | | 0 | Single Cell FISH/IF |
| H28412 | 122D | F | T21 | 2+ with patchy alveolar edema | Single Cell FISH/IF |
| H28817 | 110D | F | | 0 | FISH/IF |
| H28740 | 108D | F | T21 | 1+ with patchy alveolar edema | FISH/IF |
| H28845 | 127D | F | | 0 | FISH/IF |
| H28811 | 125D | F | T21 | 1+ | FISH/IF |

Table of fetal lungs used providing gestational age, sex, genetic anomalies, airway anomalies (Distal bronchiolar/acinar tubule dilatation), and in which assays were used. Samples used for FISH/IF are separated by a red line and are listed in their respective age and sex matched pairs of non-T21 and T21. Histopathologic analyses were performed on H&E stained sections of T21 lungs and compared to age and sex matched non-T21 controls. Scoring is classified as follows: 0 Normal, 1+ Mild, 2+ Moderate, and 3+ Severe.

Table S2. List of FISH ACD Probes

| Probe | Channel | Reference | Lot |
|---|---|---|---|
| Hs-ABCA3 | 1 | 555501 | 22164A |
| Hs-BST2 | 1 | 418651 | 22137A |
| Hs-COL1A1 | 2 | 401891 | 22137A |
| Hs-COL6A1-No-XMm | 1 | 482461 | 22137A |
| HS-COL6A2-No-XMm | 3 | 482611 | 22137A |
| Hs-FN1 | 1 | 310311 | 21355A |
| Hs-ICAM1 | 1 | 402951 | 22028A |
| Hs-IFI27 | 2 | 440111 | 22137A |
| HS-JAM2 | 1 | 412721 | 22137A |
| Hs-LYVE1 | 3 | 426911 | 22137A |
| Hs-SFTPB | 1 | 544251 | 22137A |
| Hs-SFTPC | 3 | 452561 | 22137A |
| Hs-SFTPD | 3 | 1049961 | 22137A |

Table S3. List of Antibodies

| **Primary Antibody** | **Company** | **Species** | **Catalog Number** | **Lot** | **Dilution** |
|---|---|---|---|---|---|
| CD31 | Thermo Scientific/Neomarkers | Rabbit | RB-10333-P 1 | 10333P 2107C | 1:200 |
| CDH1/E-Cadherin | BD Biosciences | Mouse | 610182 | 8274692 | 1:200 |
| ACTA2 | Dako | Mouse | M0851 | 00062068 | 1:200 |
| **Secondary Antibody** | **Company** | | **Catalog Number** | **Lot** | **Dilution** |
| Goat ∝ Mouse Alexa647 | Jackson Immunoresearch | | 115-606-146 | 155382 | 1:500 |
| Goat ∝ Rabbit Alexa647 | Jackson Immunoresearch | | 111-606-144 | 158196 | 1:500 |
| Opal 570 (FISH) | Akoya | | OP-001003 | 20213431 | 1:500 |
| Opal 520 (FISH) | Akoya | | OP-001001 | 20213026 | 1:500 |

Table S4: Epithelial Cluster Marker Genes
Table S5: Epithelial Cluster Differentially Expressed Genes
Table S6: Mesenchymal Cluster Marker Genes
Table S7: Mesenchymal Cluster Differentially Expressed Genes
Table S8: Endothelial Cluster Marker Genes
Table S9: Endothelial Cluster Differentially Expressed Genes
Table S10: Immune Cell Cluster Marker Genes

Tables S4-S10 have been uploaded as separate individual supplemental files.

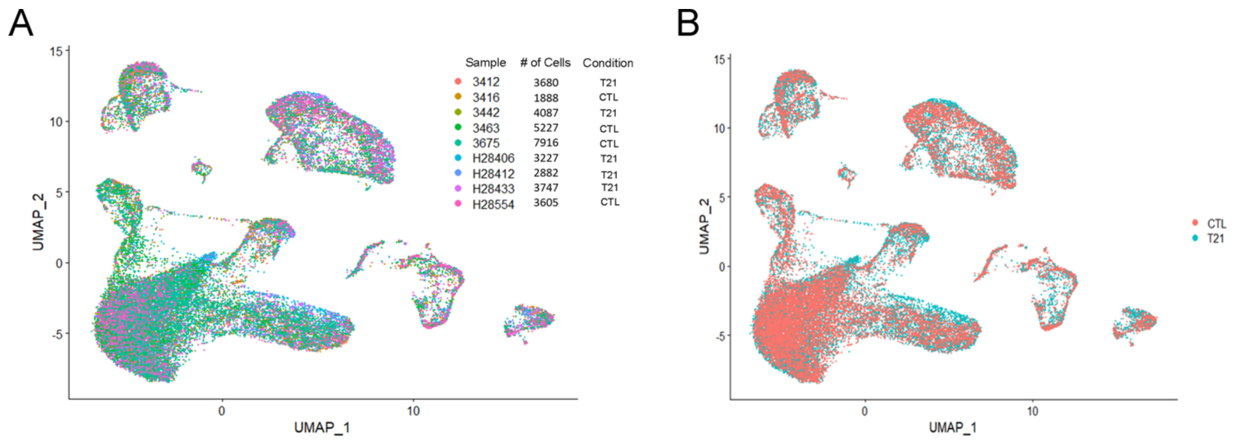


**Figure S1. T21 lungs display the same populations as observed in control non-T21 lungs.** A) UMAP displaying cell distribution by sample demonstrates uniform distribution amongst the different lung samples indicating no sample level bias. B) UMAP displaying cell distribution by condition (CTL non-T21, T21) demonstrating no unique populations in T21 lungs as compared to controls.

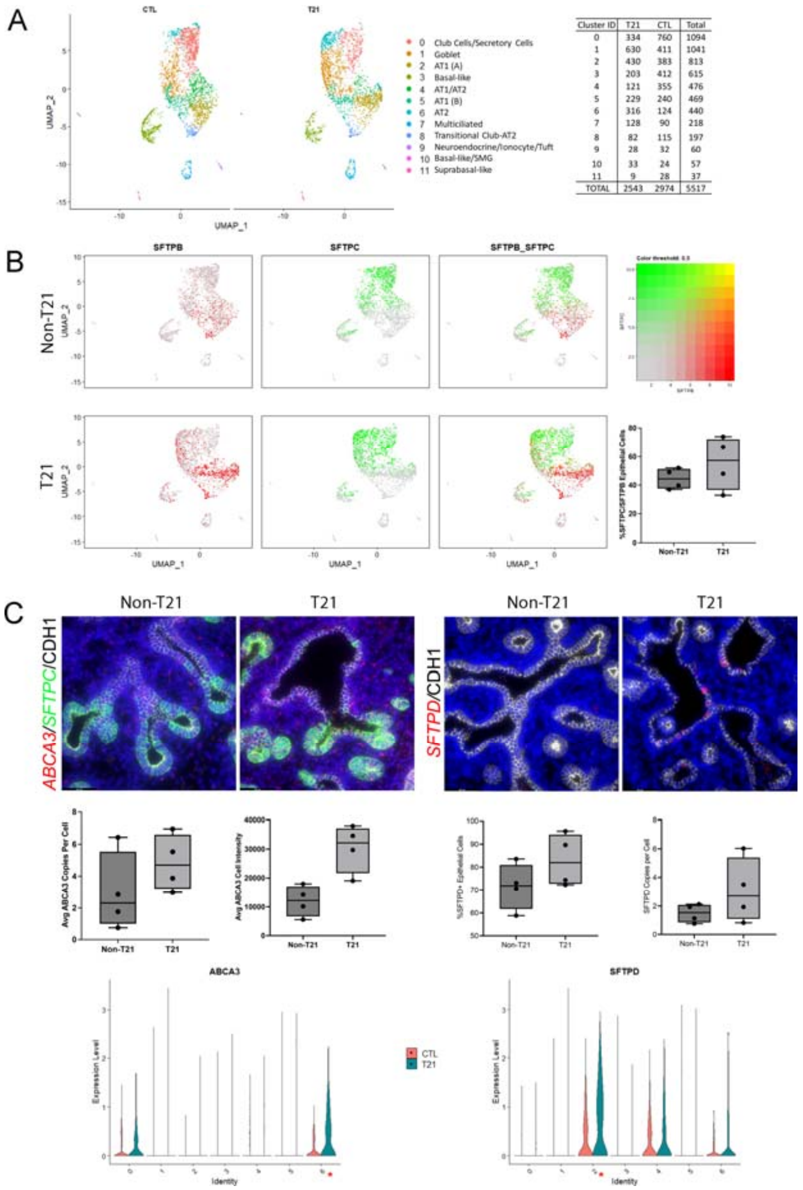


**Figure S2. Further evidence suggesting advanced differentiation in the prenatal T21 epithelium.** A) Distribution of epithelial cells by condition (non-T21 vs T21) in UMAP shows no unique epithelial cell clusters in the fetal T21 lung. Table shows cell distribution by condition for each sub-cluster. B) Assessing epithelial co-expression of *SFTPB* (Red) and *SFTPC* (Green) by condition and represented by UMAP. Each dot represents a single cell. Cells demonstrating *SFTPB/SFTPC* co-expression is represented in yellow. Single cell RNAseq data show there are more *SFTPB/SFTPC* co-expressing cells in the fetal T21 lung as compared to non-T21, which was further suggested by a trend observed in FISH quantification (n=4, p=0.1532). C) FISH of non-T21 control and T21 age and sex matched fetal lungs demonstrate a trend toward increased expression for *ABCA3*. Although the percentage of epithelial cells positive for *ABCA3* is comparable between T21 and non-T21 lungs, there appears to be on average more transcripts per cell (p=0.1310) in the T21 lungs with a stronger intensity (p=0.0669). Furthermore, our single cell data demonstrates that *ABCA3* is differentially expressed in both Cluster 0 (Club Cells/Secretory Cells) and Cluster 6 (AT2), as represented by violin plot. FISH of non-T21 control and T21 age and sex matched fetal lungs also demonstrate a trend toward increased expression for *SFTPD*. The percentage of epithelial cells positive for *SFTPD* (p=0.1985) as wells as the number of *SFTPD* transcripts per cell (p=0.1941) are trending towards an increase in the T21 lungs as compared to the non-T21. Furthermore, our single cell data demonstrates that *SFTPD* is differentially expressed in several clusters, such as Cluster 2 (AT1 (A)), Cluster 4 (AT1/AT2) and Cluster 6 (AT2), as represented by violin plot. (N=4 for each condition; Scale bar: 64.7um). Box Plots: center line, mean; box limits, upper and lower quartiles; whiskers, minimum to maximum

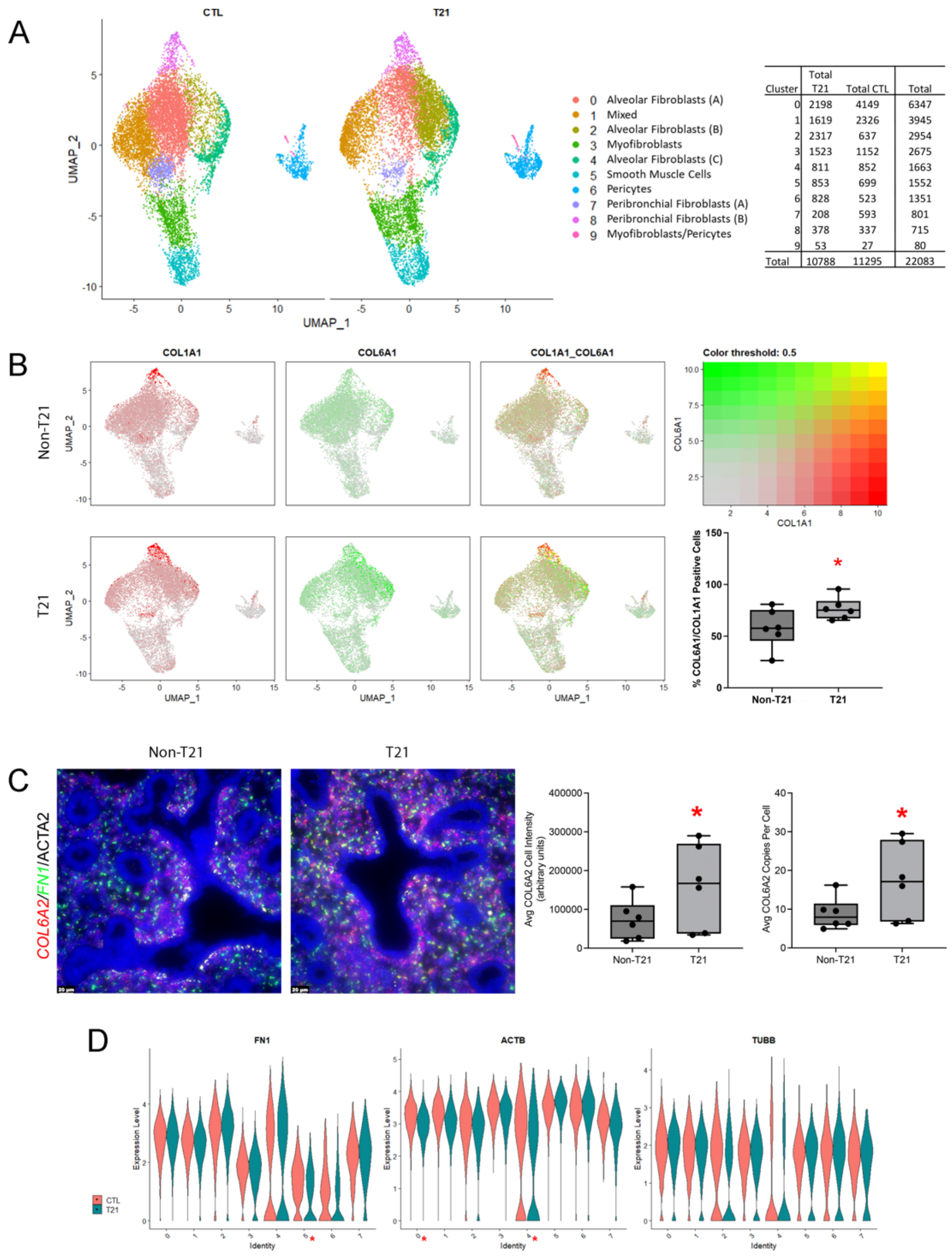


**Figure S3. ECM components are significantly upregulated in prenatal T21 mesenchyme.** A) Distribution of mesenchymal cells by condition (non-T21 vs T21) in UMAP shows no unique mesenchymal cell clusters in the fetal T21 lung. Table shows cell distribution by condition for each sub-cluster. B) Assessing mesenchymal co-expression of *COL1A1* (Red) and *COL6A1* (Green) by condition and represented by UMAP. Each dot represents a single cell. Cells demonstrating *COL1A1/COL6A1* co-expression is represented in yellow. Single cell RNAseq data show there are more *COL1A1/COL6A1* co-expressing cells in the fetal T21 lung as compared to non-T21, which was further confirmed in FISH quantification (n=6, p=0.0289). C) FISH of non-T21 control and T21 age and sex matched fetal lungs demonstrate significantly increased expression of *COL6A2* in both the copy number per cell as well as signal intensity in T21 lungs as compared to controls (COL6A2: Copy number p=0.0346, Intensity p=0.0322, n=6). D) Violin plots demonstrating differential expression of mesenchymal genes that have been shown to be important branching and smooth muscle contraction: *FN1, ACTB*, and *TUBB*. Significance (p<0.05) is indicated by * next to cluster number on x-axis. Box Plots: center line, mean; box limits, upper and lower quartiles; whiskers, minimum to maximum

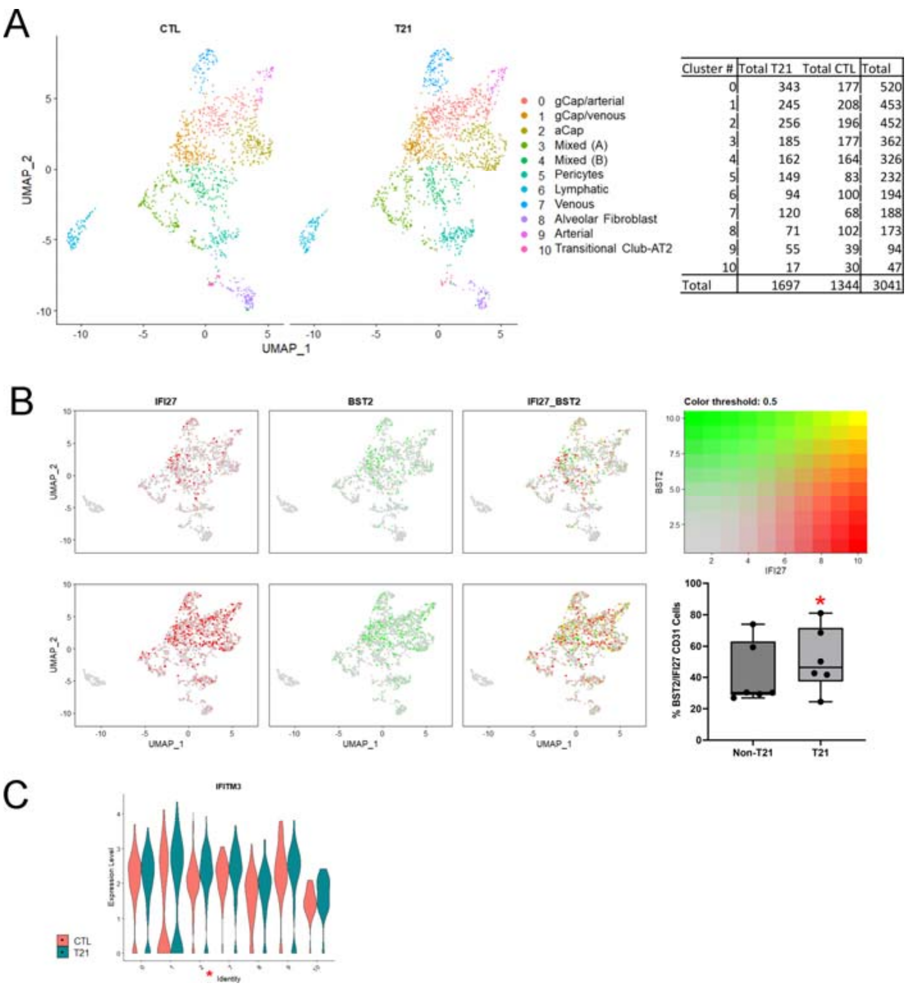


**Figure S4. Type I IFN targets are upregulated in prenatal T21 endothelium.** A) Distribution of endothelial cells by condition (non-T21 vs T21) in UMAP shows no unique endothelial cell clusters in the fetal T21 lung. Table shows cell distribution by condition for each sub-cluster. B) Assessing endothelial co-expression of type I IFN downstream targets *IFI27* (Red) and *BST2* (Green) by condition and represented by UMAP. Each dot represents a single cell. Cells demonstrating *IFI27/BST2* co-expression is represented in yellow. Single cell RNAseq data show there are more *IFI27/BST2* co-expressing cells in the fetal T21 lung as compared to non-T21, which was further confirmed in FISH quantification (n=6, p=0.0230). C) Violin plot of *IFITM3*, another downstream target of Type I IFN signaling, shows that it is differentially upregulated in several of the T21 endothelial cell clusters. Significance ($p<0.05$) is indicated by * next to cluster number on x-axis. Box Plots: center line, mean; box limits, upper and lower quartiles; whiskers, minimum to maximum

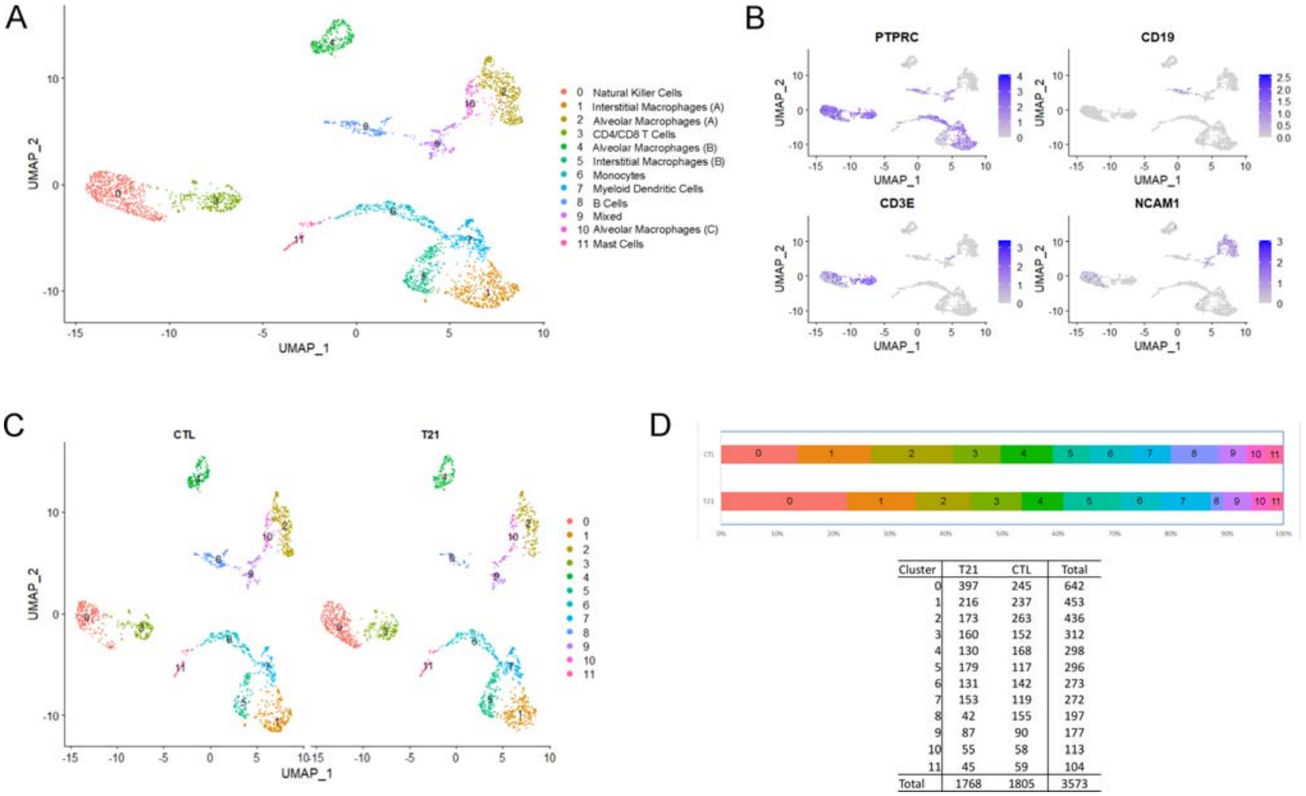


| Cluster | T21 | CTL | Total |
|---|---|---|---|
| 0 | 397 | 245 | 642 |
| 1 | 216 | 237 | 453 |
| 2 | 173 | 263 | 436 |
| 3 | 160 | 152 | 312 |
| 4 | 130 | 168 | 298 |
| 5 | 179 | 117 | 296 |
| 6 | 131 | 142 | 273 |
| 7 | 153 | 119 | 272 |
| 8 | 42 | 155 | 197 |
| 9 | 87 | 90 | 177 |
| 10 | 55 | 58 | 113 |
| 11 | 45 | 59 | 104 |
| Total | 1768 | 1805 | 3573 |

**Figure S5. Immune cell lineage shows the fewest differences between the T21 and non-T21 lungs**. A) Uniform Manifold Approximation and Projection (UMAP) displays the 12 unique immune cell sub-clusters. Each dot represents a single cell and individual clusters are colored and annotated based on cell type associations identified by ToppFun, using marker genes for each individual cluster. B) Gene expression patterns for individual canonical immune cell markers (*PTPRC, CD19, CD3E, NCAM1*) overlaid on UMAP plots confirming immune cell lineage. Each point represents a single cell, with blue color indicating expression level of the specified marker gene (darker shade is higher expression). C) Distribution of immune cells by condition (non-T21 vs T21) in UMAP shows no unique immune cell clusters in the fetal T21 lung. D) Proportional differences in immune cell sub-clusters between non-T21 and T21 cohorts show that none of the clusters are significantly different. Table shows cell distribution by condition for each sub-cluster.
